# Fast and accurate approximation of the joint site frequency spectrum of multiple populations

**DOI:** 10.1101/2020.05.01.073213

**Authors:** Ethan M. Jewett

## Abstract

The site frequency spectrum (SFS) is a statistic that summarizes the distribution of derived allele frequencies in a sample of DNA sequences. The SFS provides useful information about genetic variation within and among populations and it can used to make population genetic inferences. Methods for computing the SFS based on the diffusion approximation are computationally efficient when computing all terms of the SFS simultaneously and they can handle complicated demographic scenarios. However, in practice it is sometimes only necessary to compute a subset of terms of the SFS, in which case coalescent-based methods can achieve greater computational efficiency. Here, we present simple and accurate approximate formulas for the expected joint SFS for multiple populations connected by migration. Compared with existing exact approaches, our approximate formulas greatly reduce the complexity of computing each entry of the SFS and have simple forms. The computational complexity of our method depends on the index of the entry to be computed, rather than on the sample size, and the accuracy of our approximation improves as the sample size increases.

## 1. Introduction

The site frequency spectrum (SFS) is a statistic that records the distribution of allele frequencies across one or more populations (Hartl and Clark 2007, Wakeley 2008). The distribution of allele frequencies contains information about the size history of a population and evolutionary factors such selective pressures (Watterson 1975, McCoy et al. 2014, Bhaskar and Song 2014, Tajima 1989, Fay and Wu 2000, Nielsen et al. 2005), making the SFS is a useful statistic for performing inference under population-genetic models (Marth et al. 2004, Keinan et al. 2007, Gutenkunst et al. 2009, Excoffier et al. 2013).

Many methods have been developed for computing the SFS among populations of time-varying size connected by migration. Methods based on the Wright-Fisher diffusion model (Gutenkunst et al. 2009, Gravel et al. 2011, Lukić and Hey 2012) are typically quite fast when the number of populations is small. However, for multiple populations, the SFS is a multidimensional array in which the number of dimensions equals the number of populations. Because diffusion-based methods must keep track of the full multidimensional distribution of allele frequencies when integrating forward in time, these approaches rapidly become computationally challenging as the number of populations increases.

In contrast to diffusion-based approaches, methods based on the coalescent model only consider the histories of alleles that are observed in the present-day sample. Thus, coalescent approaches allow the computation of the SFS term-by-term. Such approaches can lead to considerable improvements in both speed and accuracy for computing subsets of entries of the SFS, which can improve the efficiency of inference. Coalescent formulas for computing the expected SFS have been obtained for both single populations of time-varying size and for multiple populations connected by migration (Wakeley and Hey 1997, Chen 2012, Chen and Chen 2013). Recently, Kamm et al. (2017) greatly improved the efficiency and numerical stability of coalescent approaches by developing a recursive algorithm for computing the SFS in populations of time-varying size with pulse migrations among them. Jouganous et al. (2017) have also developed a method that is a variation on diffusion approaches, which allows the SFS to be computed for larger numbers of populations.

Although the methods of Kamm et al. (2017) and Jouganous et al. (2017) have made computation of the SFS extremely fast, it may be possible to reduce the complexity of computing terms of the SFS still further. A potential approach for reducing the complexity of coalescent methods is to derive accurate approximations of the SFS using deterministic approximations of the number of ancestral coalescent lineages remaining in the population at each time in the past (Griffiths 1984, Slatkin and Rannala 1997, Volz et al. 2009, Maruvka et al. 2011, Chen and Chen 2013, Jewett and Rosenberg 2014). These approximations can be used to derive accurate approximate formulas for a variety of useful population genetic quantities (Maruvka et al. 2011, Chen and Chen 2013, Jewett and Rosenberg 2014) and they can be computed under complicated demographic histories that are difficult for classical coalescent models (Jewett and Rosenberg 2014).

Chen and Chen (2013) derived an approximation of the single-population SFS using a formula by Polanski and Kimmel (2003), in which the single-population SFS is expressed as a sum over expected coalescent waiting times. The approximate formulas of Chen and Chen (2013), which were obtained by replacing exact expressions for expected coalescent waiting times in the Polanski and Kimmel formula with accurate approximations, are fast and accurate for computing the SFS in a single population.

Here, we take an analogous approach to that of Chen and Chen (2013) to compute an accurate approximation of the SFS in a set of populations with migration. However, rather than using the expression from Polanski and Kimmel (2003) for the SFS as a sum over expected waiting times, we take a new approach, deriving formulas for the SFS under an approximate coalescent framework in which the number of lineages as a function of time is deterministically equal to its expected value. Using this approach, we show that the SFS can be expressed as a sum over the expected total length of branches ancestral to subsets of sampled sequences. As we will see, this alternative formulation of the SFS provides a simple and intuitive way of deriving approximations of the SFS in multiple populations connected by migration. This approach allows us to obtain formulas for the SFS that have simple expressions and low computational complexity.

## 2. A summary of the main results

In a single population, the SFS for a sample of *n* sequences is an ordered tuple ***ξ***_*n*_ = (*ξ*_*n*,1_, …, *ξ*_*n,n*−1_) of length *n* − 1 in which the *i*th term records the number of polymorphic sites at which the derived allele appears in exactly *i* out of *n* sequences. For *P* populations with *n*_*p*_ sequences sampled from population *p*, the SFS is a *P*-dimensional array in which entry *i*_1_, …, *i*_*P*_ records the number of polymorphic sites at which the derived allele appears in *i*_*p*_ copies in population *p*, for *p* = 1, …, *P*. We first present approximate formulas for the SFS in the case of a single population of piecewise constant size and then consider more complicated models. All derivations are deferred to Section 3.

### 2.1. Computing the SFS in a single population

Consider a single population of piecewise constant size like the one shown in the figure in Box 1. In such a population, the size *N*(*t*) at time *t* changes over the course of *K* different time intervals 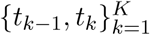 satisfying *t*_0_ < *t*_1_ < … < *t*_*K*_ ≤ ∞ in such a way that *N*(*t*) = *N*(*t* _*k*−1_) for *t* ∈ [*t* _*k*−1_, *t* _*k*_). We denote the relative population size at time *t* by *ν*(*t*) = *N*(*t*)/ *N* for some reference effective population size *N*, where time is measured in units of 2*N* generations.

#### Box 1

Computing 𝔼***ξ***_*n*_ in a single population of piecewise constant size.

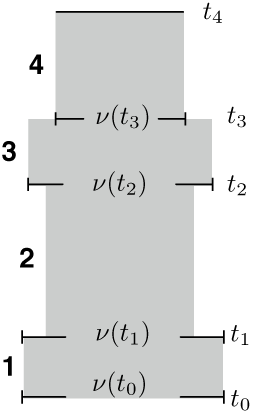

Consider a population size history defined piecewise over *K* different time intervals 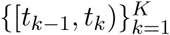 for times satisfying *t*_0_ < *t*_1_ < < *t*_*K*_≤ ∞, where time is measured in coalescent units of 2*N* generations with respect to a reference population of arbitrary size *N* diploids. Suppose that the relative population size in the kth time interval is given by *ν*(*t*) = *ν*(*t*_*k*−1_) for *t* ∈ [*t*_*k*−1_, *t*_*k*_). An example of such a population history with four time intervals is shown on the left.

The exact expectation of the ith entry of the classical SFS ***ξ***_*n*_ is

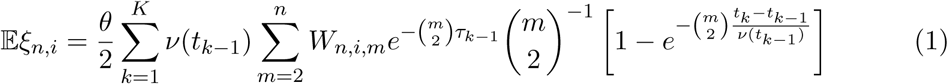

where

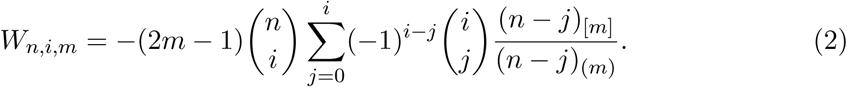

The quantity *W*_*n,i,m*_ can also be computed efficiently using the following recursion from Kamm et al. (2017)

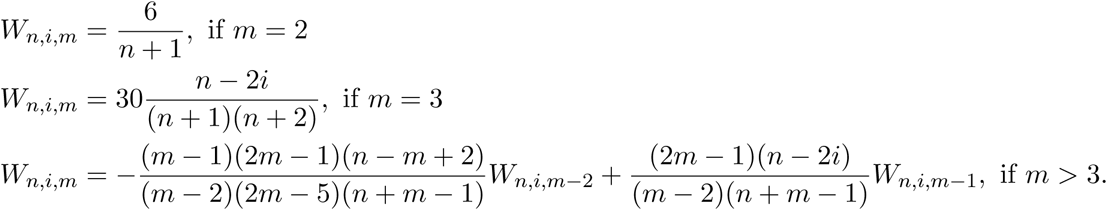

**An approximate formula:** The expectation of the ith term of the SFS can be approximated by approximated by

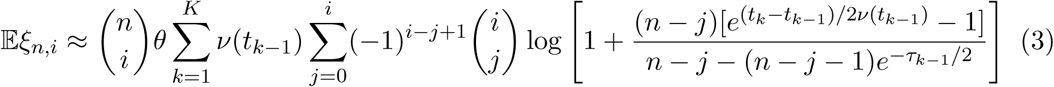

whenever *t*_*K*_ < ∞, where 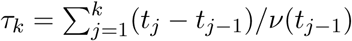 and 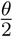 is the mutation rate in the coalescent.

In Section 3.3, we show that our approach yields the exact formula for the SFS given in Equation (1), which is equivalent to Equation (10) of Kamm et al. (2017). Because the formula appears in a general form expressed as an integral in Kamm et al. (2017), we report the specific simple form for a piecewise constant population here for completeness. Note that when *t*_*K*_ < ∞, Equation (1) computes the truncated SFS derived in Kamm et al. (2017). In Section 3.4, we also show that the expectation of ***ξ***_*n*_ in a population of piecewise constant size can be approximated using Equation (3) in Box 1.

The approximate formula for the SFS in Box 1 provides a fast method for computing the SFS that is simple to implement. Although diffusion approaches are still the fastest methods for computing the complete SFS for large *n*, Equation (3) provides an improvement in efficiency over existing methods for computing subsets of entries. In particular, the complexity of Equation (3) does not depend on the sample size *n*. Instead, it depends linearly on the index *i* of the computed term. The complexity for computing the first L terms of the SFS is *O*(*L*^2^*K*), where *K* is the number of piecewise constant epochs, which is lower than the *O*(*LnK*) complexity of Equation (1), which is the current state of the art.

Panels A and E of Figure 1 show the runtimes of evaluating the formulas in Box 1 implemented in Mathematica, compared with those of the SFS software packages *momi* and *∂a∂i*. For all plots in Figure 1, the SFS was computed for a population with a bottleneck. The population size history is given by *ν*(*t* _0_) = 1, *ν*(*t*_1_) = 0.5, *ν*(*t* _2_) = 1 at times *t* _0_ = 0, *t* _1_ = 0.1, *t*_2_ = 0.2, and *t* _3_ = ∞. When computing Equation (3), we require *t* _3_ < ∞ so we chose the large value *t* _3_ = 20. As expected, the runtime of the approximate formula (Equation 3) is constant in *n*, whereas the runtimes of *momi* and *∂a∂i* increase with *n*.

**Figure 1.**
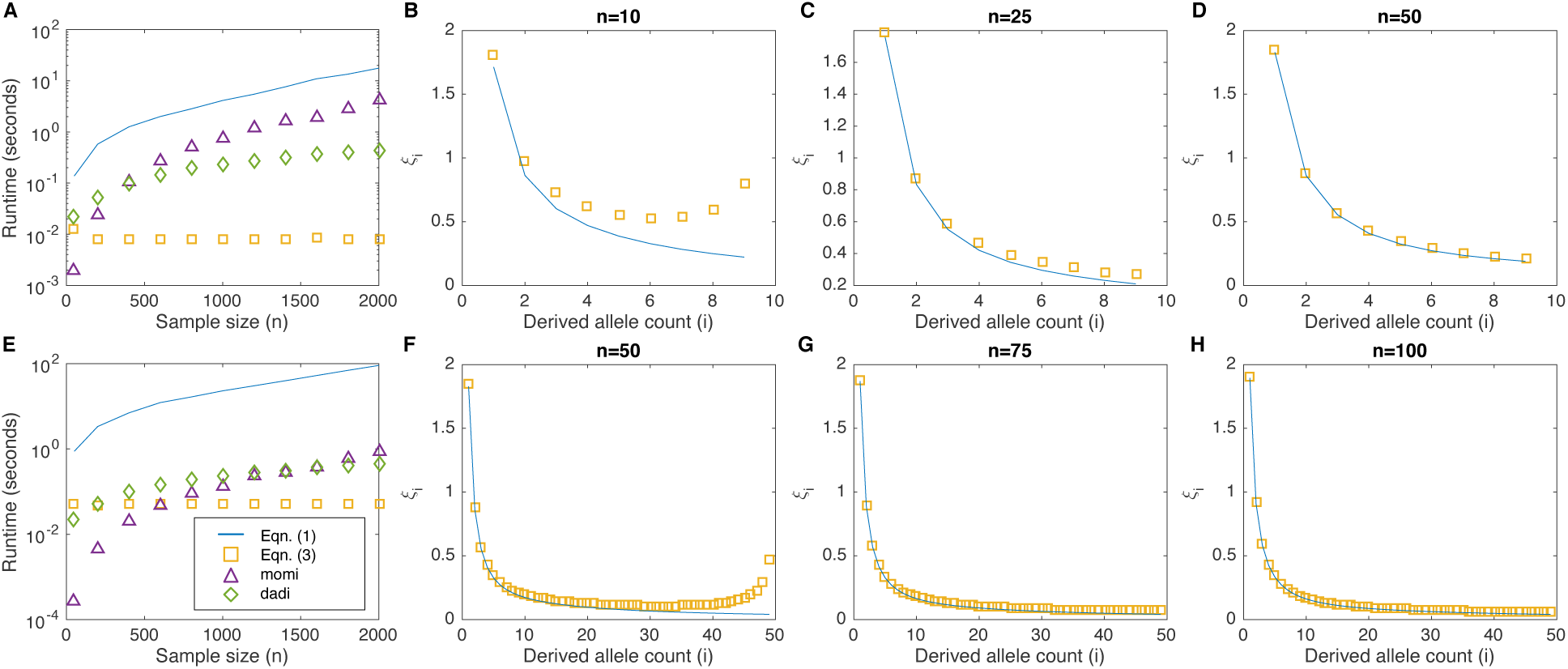
Accuracy and timing of the formulas in Box 1 for a population with a bottleneck. All panels correspond to a population history in which the size is given by *ν*(*t*_0_) = *ν*(*t*_2_) = 1, *ν*(*t*_1_) = 0.5, at times *t*_0_ = 0, *t*_1_ = 0.1, and *t*_3_ = 0.2. Panel A shows the runtime for computing the top 9 entries of the SFS for different sample sizes. Panels B-D show the accuracy in the top 9 entries of the SFS as the sample size *n* is increased from 10 to 50. Panel E shows the runtime for computing the top 49 entries, and panels F-H show the accuracy in the top 49 entries as the sample size *n* is increased from 50 to 100.

It is evident from Panels A and E of Figure 1 that, even though the asymptotic complexity of Equation (1) is the same as that of *momi*, additional speed-ups obtained in the implementation of *momi* make *momi* extremely fast for computing low-order terms of the SFS when sample sizes are small to moderate. In fact, both *momi* and *∂a∂i* are fast in the case of a single population.

## 2.2. Computing the joint SFS in multiple populations with pulse migrations

For samples taken from *P* different populations, the SFS is a *P*-dimensional array. In particular, if *n*_*p*_ homologous sequences are sampled from population *p* (*p* = 1, …, *P*), then ***ξ*** is an array with dimensions *n*_1_ × *n*_2_ × … × *n*_*P*_ in which entry (*i*_1_, …, *i*_*P*_) records the number of polymorphic sites at which the derived allele appears on *i*_1_ sequences from the first population, *i*_2_ sequences from the second population, and so on.

We show in Section 3 that the SFS for a collection of populations of piecewise constant size connected by instantaneous pulse migrations can be approximated using Equation (4) in Box 2. Figure 3 shows the accuracy and timing of Equation (4) compared with momi and ∂a∂i for the case of two populations of sizes *ν*_1_ = 3 and *ν*_2_ = 2 that split from an ancestral population of size *ν*_*A*_ = 1 at *t* = 0.05 coalescent time units in the past. From Panels A-F of Figure 3, it can be seen that the approximate formula accurately captures the lower-order terms of the SFS even when the number of samples is small (e.g., *n*_1_ = *n*_2_ ≈ 20).

**Figure 2.**
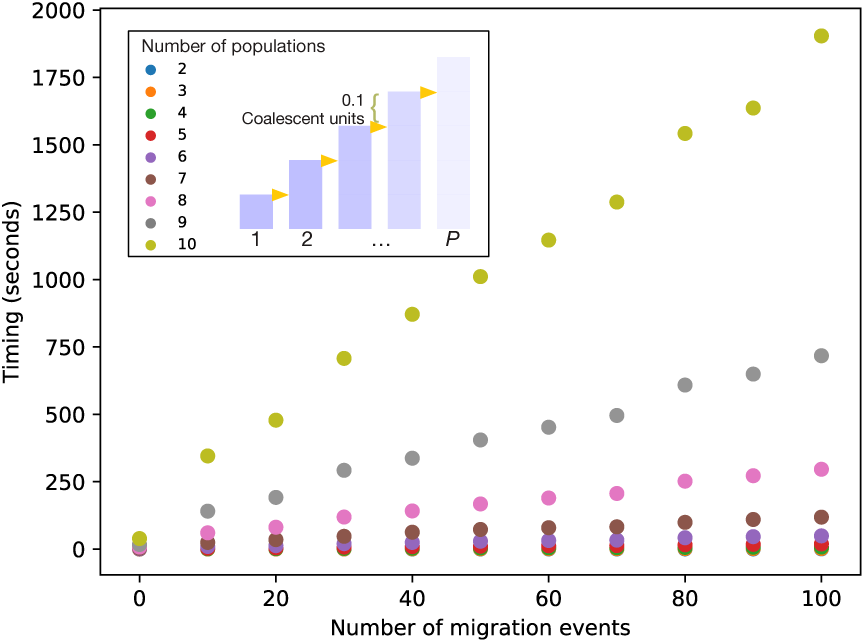
Time for computing the first term of the SFS using the approximation in Box 2, for different numbers of populations and pulse migration events. The form of the population history is shown in the inset diagram. In this model, a new population branches from its ancestral population every 0.1 coalescent units and each population has constant size *N* = 10, 000. Migrations from the descendant population to the ancestral population are evenly spaced between the present-day and the founding event and 10% of the lineages in the descendant population migrate in each event. The sample size from each population is 50 lineages.

**Figure 3.**
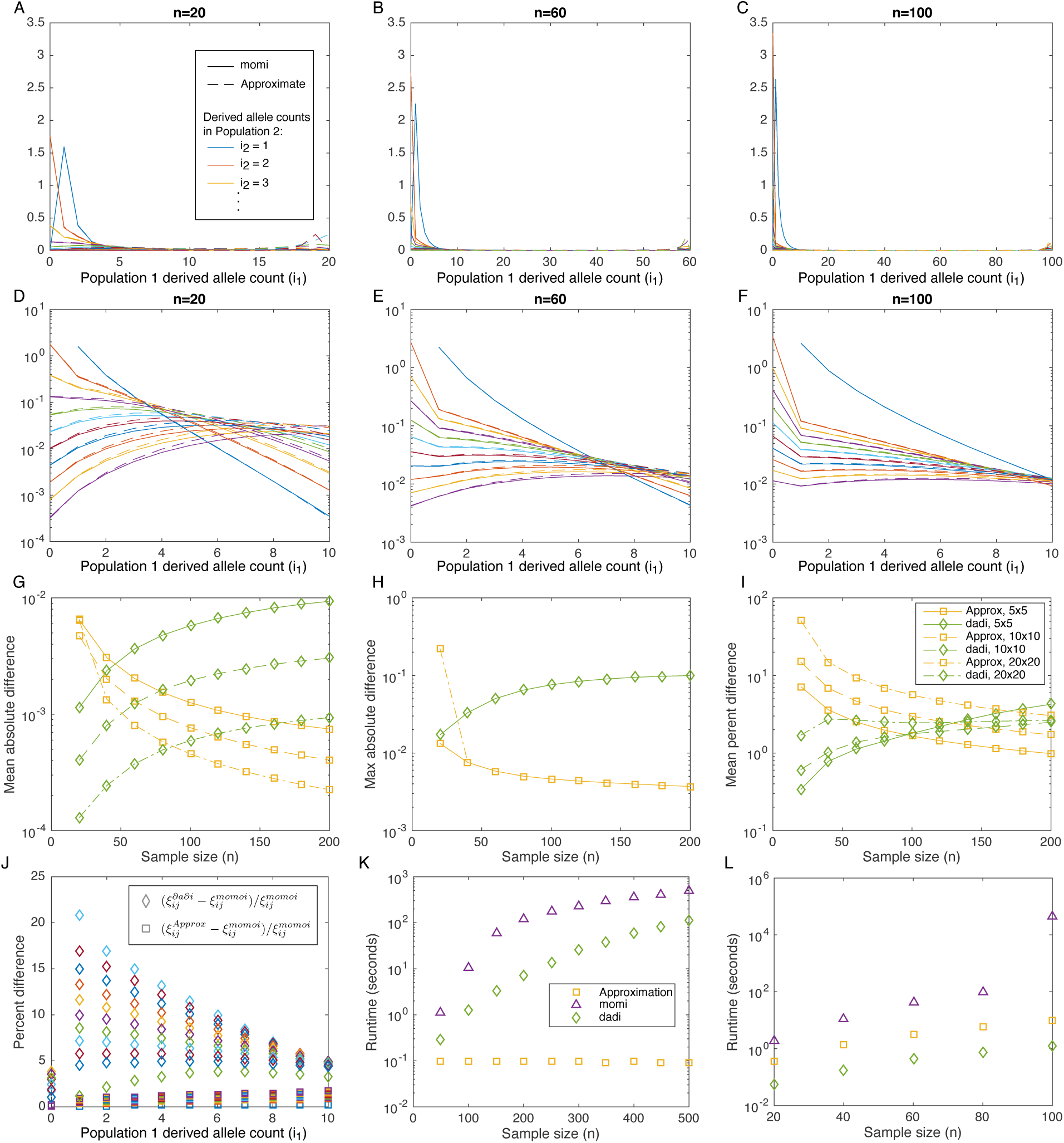
Accuracy and timing of Equation (4) in Box 2 for two populations of constant sizes *ν*_1_ = 3 and *ν*_2_ = 2 that diverged at time *t* = 0.05 from a population of size *ν*_*A*_ = 1. No migration occurred after the split. Panels A, B, and C show a comparison between the approximate SFS (Equation 4) and the SFS generated by *momi*. Panels D-F show the same comparison, zoomed-in on the first 11 × 11 corner of the SFS corresponding to counts *i* _1_ = 0, …, 10 in population 1 and *i*_2_ = 0, …, 10 in population 2. Panels *G* through *I* show average errors over subsets of SFS terms. In Panels G-I, *∂a∂i* and Equation (4) are separately compared to *momi*, which is taken as the true SFS. Averages are taken over subsets of SFS terms corresponding to the first 6 × 6 terms, the first 11 × 11 terms, and the first 21 × 21 terms. Panel *J* shows the percent error in each of the top 11 × 11 terms for a sample size of *n*_1_ = *n* _2_ = *n* = 500. Panel K shows the runtime for computing the first 11 × 11 corner of the SFS. Panel L shows the runtime for computing the full spectrum. Colors are consistent among panels A-F and J, and separately among panels G, H, I, K, and L. Plot marker shapes are consistent across panels.

### Box 2

Approximating 𝔼*ξ*_n_ in *P* piecewise constant populations with pulse migrations.

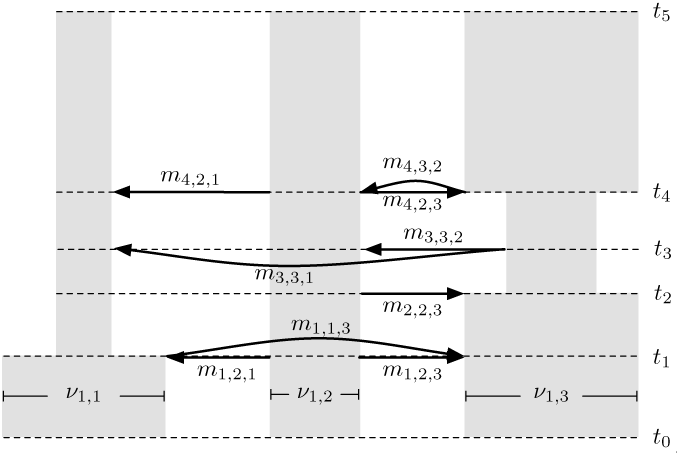

Consider a collection of*P* piecewise constant populations with pulse migrations in which the relative size of population *p* is *ν*_*p*_(*t*) = *ν*_*k,p*_ for *t* ∈ [*t*_*k*−1_, *t*_*k*_) measured in units of 2*N* generations and satisfying *t*_0_ < *t*_1_ < < *t*_*K*_ < ∞. Let *m*_*k,p,p*′_ be the fraction of population *p*′ that instantaneously immigrates from population *p* at the top of interval *k*, looking forward time and let ***M***_*k*_ be the *P* × *P* matrix whose (*p, p*′) entry is *m*_*k,p,p*′_ if *p* ≠ *p*′ and 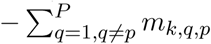 if *p* = *p*′. Note that a population split at the top of interval k is handled by setting *m*_*k,p,p*′_ = 1 so that all lineages migrate into one population. Suppose that *n*_*p*_ sequences are sampled from population *p*, for *p* = 1, …, *P*, and let **n** = (*n*_1_, …, *n* _*P*_). Then entry (*i*_1_, …, *i*_*P*_) of the SFS ***ξ***_**n**_ can be computed using the formula

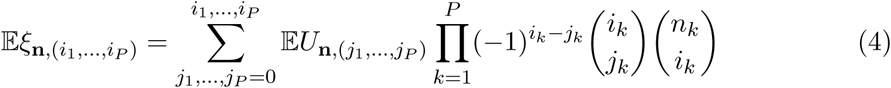

where

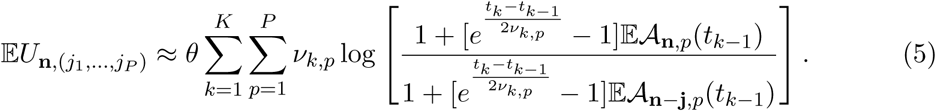

In Equation (4), the quantity 𝔼*𝒜*_**i**,*p*_(*t*_*k*_) is the expected number of ancestral lineages at time *t*_*k*_ in population *p*, before migration occurs looking forward in time, given that ***i*** = (*i*_1_, …, *i* _*P*_) lineages are initially sampled at time zero. The quantity *𝒜*_**i**,*p*_(*t*_*k*_) can be found recursively using

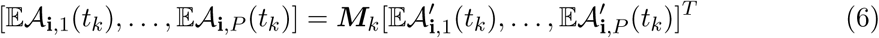

where

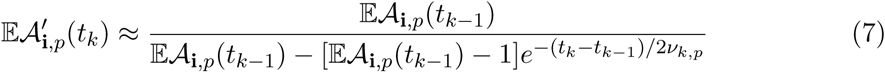

is the number of lineages remaining in population *p* immediately after migration occurs at time *t*_*k*_, looking forward in time, and 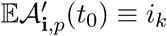.

There is always a certain amount of error in the higher order terms of the SFS, which is visible as the right-hand-side of Panels A-C of Figure 3. However, for lower order terms of the SFS, the approximation becomes more accurate than *∂a∂i* as the sample size increases. This can be seen in panels G, H and I of Figure 3, which compare errors in the approximation with errors in *∂a∂i*, taking the values of *momi* as the truth. Panels G and H show average absolute errors, 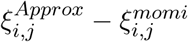 and 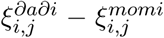, whereas Panel I shows average percent errors, 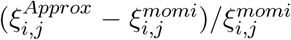 and 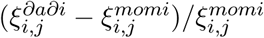.

As expected, the absolute error in the approximation is greatest in the terms of greatest magnitude, which are typically the low order terms of the SFS. This can be seen in Figure 3G, which shows the mean absolute difference between the approximation and *momi* for different subsets of SFS terms. In particular, the yellow solid line shows the average absolute error across the first 5 × 5 terms of the SFS, whereas the dashed dotted yellow line shows the average absolute error across the expanded set containing the first 20 × 20 terms. Because the solid line in Figure 3G is above the dashed and dashed-dotted lines, it can be seen that the average absolute error is greater for the first 5 × 5 terms than for the higher order terms.

The fact that the low-order (high magnitude) terms of the SFS have the greatest absolute error is supported by figure Figure 3H, which shows that the maximum absolute error is the same when considering the first 5 × 5 terms, the first 10 × 10 terms, or the first 20 × 20 terms (the solid, dashed, and dashed-dotted lines are on top of one another), indicating that the maximum absolute error is in the set of first 5 × 5 term. In contrast, the mean relative error increases as we consider higher order terms of the SFS (Figure 3I), suggesting that the greatest relative error is in the higher order terms.

In Figures 3G-H, we have also compared *∂a∂i* to *momi* (green lines). From Figures 3G-H it can be seen that the diffusion approximation in *∂a∂i* diverges from the values computed by *momi* as the sample size increases.

When computing the low-order entries of the SFS in the case of multiple populations, the approximate SFS is also faster than existing approaches. Figure 3K shows the computation time of the first 11 × 11 entries of the SFS corresponding to counts counts *i*_1_ = 0, …, 10 in population 1 and *i*_2_ = 0, …, 10 in population 2. From Figure 3K, it can be seen that the approximation can be computed in constant time in the sample size n, making it faster than the current state of-the-art approaches for computing terms of the SFS when the sample size is large. Although the approximation is the fastest approach for computing each individual SFS term, Figure 3L suggests that *∂a∂i* is still the fastest method for computing the full spectrum.

Figure 2 shows the computation time for the approximation under increasingly complicated demographic histories with increasing numbers of populations and pulse migration events. In particular, we modeled multiple populations by considering a serial founder model in which each population gave rise to an offspring population via a pulse migration consisting of 10% of its lineages. To evaluate the computation time of the approximation for *p* populations, we considered the youngest *p* populations. To evaluate the computation time in the presence of pulse migrations, we we added increasing numbers of pulse migrations between each population and its parent population, evenly spaced across the age of the younger population. The computation time grows quadratically in the number of populations and linearly in the number of pulse migration events (Figure 2).

## 3. Derivation of the formulas

In this section we derive the equations presented in Sections 2.1 and 2.2. Our approach is to first express the SFS as a sum of expected total branch lengths ancestral to different subsets of sampled sequences. This approach is analogous to that of Polanski and Kimmel (2003) who expressed the SFS as a sum of expected coalescence waiting times. As we will see, the exact formulas for the SFS obtained by the two approaches are the same; however, our approach of integrating over branch lengths makes it straightforward to derive fast approximate formulas in the case of multiple populations.

To facilitate the derivations, we define notation that we will use throughout this section. For a set of *n* sequences sampled from a single population, let *S*_*i*_ denote a particular subsample of *i* sequences. Let *π*(*S*_*i*_) denote the number of sites at which the derived allele is private to the sample *S*_*i*_ and let *α*(*S*_*i*_) denote the number of sites at which the derived allele is ancestral to all of the *i* sequences in the sample, and to no others. The quantities *π*(*S*_*i*_) and *α*(*S*_*i*_) are illustrated in Figure 4 for a subsample of size *i* = 3 sequences at 3 different loci.

**Figure 4.**
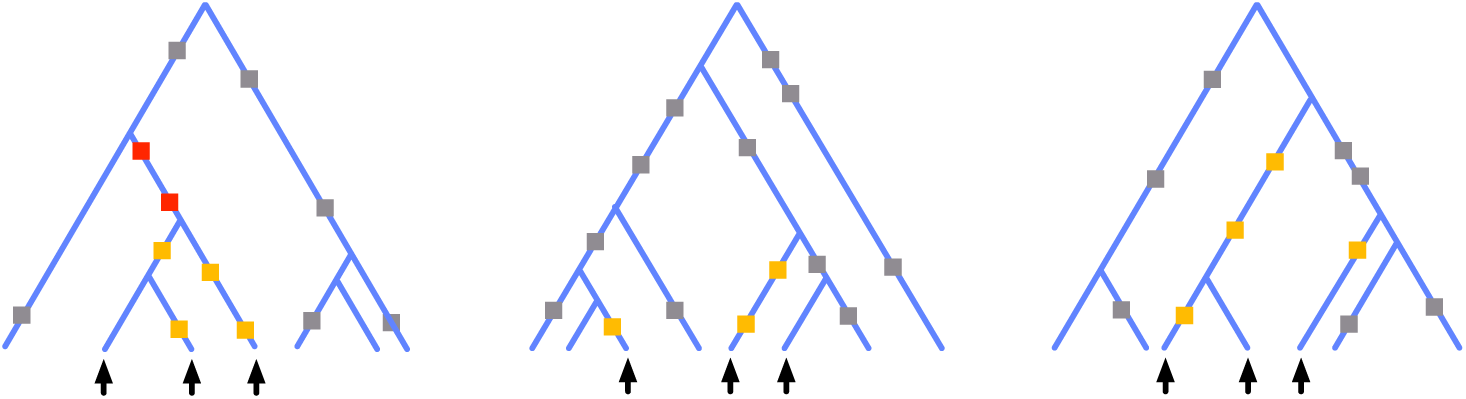
The quantities *π*(*S*_*i*_) and *α*(*S*_*i*_). A subsample of *i* = 3 sequences (arrows) is shown for a total sample of *n* = 7 sequences. Trees are shown for three different loci along these sequences. Squares indicate mutations. For the indicated subsample, the number of private sites is *π*(*S*_*i*_) = 13 (sum of all red and yellow squares), whereas the number of ancestral sites is *α*(*S*_*i*_) = 2 (red squares). Note that private sites are counted even when the subsample does not form a monophyletic clade.

The quantity *α*(*S*_*i*_) is closely related to the ith term of the ***ξ***, which is simply the sum of *α*(*S*_*i*_) over all subsets of size *i*; the summation yielding the total number of segregating sites that appear in exactly *i* lineages. To compute *α*(*S*_*i*_) directly, we could consider the number of lineages that are ancestral to all members of *S*_*i*_ at each time *t* in the past and then integrate this quantity over all time, multiplied by the rate of new mutations. Our approach is to observe that the quantity *π*(*S*_*i*_) is considerably easier to compute, being the integral over the total number of branches ancestral to the set *S*_*i*_. The quantity *α*(*S*_*i*_) is related to *π*(*S*_*i*_) via the principle of inclusion and exclusion. We now provide the details of this computation.

Considering all possible subsets 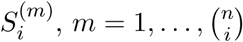, of {1, …, *n*} with exactly *i* elements, we define a quantity *U*_*n,i*_ as

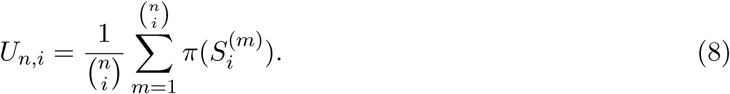

Intuitively, the quantity *U*_*n,i*_ is the average number of sites that are private to a set of *i* lineages. Equation (8) is similar in form to the expression for the ith term of the SFS, which can be expressed as

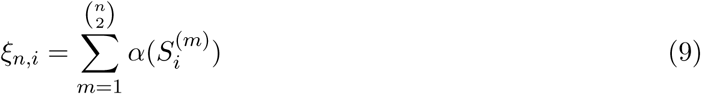

By relating *π*(*S*_*i*_) with *α*(*S*_*i*_), we can establish a relationship between 𝔼*ξ*_*n,i*_ with 𝔼*U*_*n,i*_. Because 𝔼*U*_*n,i*_ is proportional to an expected branch length, this approach allows us to establish a formula for 𝔼***ξ***_*n*_ in terms of expected sums of branch lengths. This is the approach that we will take here to derive the approximations given Section 2.

### 3.1. The relationship between *U*_*n*_ and *ξ*_*n*_ in one population

We first establish the relationship between *U*_*n,i*_ and *ξ*_*n,i*_ in a single population before proceeding to the case of multiple populations. To derive the relationship, note that a site is private to *S*_*i*_ if and only if it is ancestral to some subset 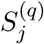 of *j* ≤ *i* sequences satisfying 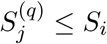, and to no others. Moreover, the set of mutations that are ancestral only to one particular subset 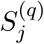 of size *j* is mutually exclusive of the mutations ancestral only to a different subset 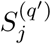 of size j. Thus, the number *π*(*S*_*i*_) of private sites in *S*_*i*_ is equal to the sum over the ancestral sites in every subset of *S*_*i*_ of every size. In other words, we have

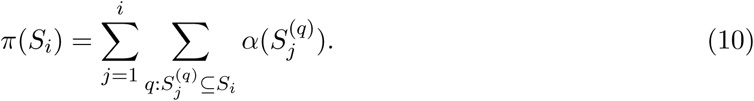

Plugging Equation (10) into the definition in Equation (8) gives

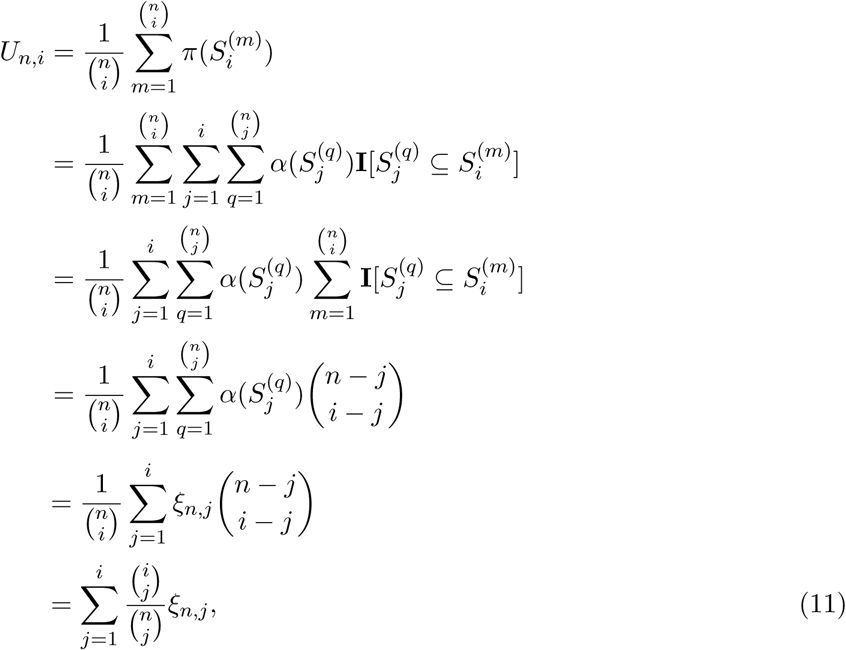

where **I**[·] is the the indicator function, which is equal to 1 when its argument is true and 0 otherwise. The second equality in Equation (11) follows by plugging Equation (10) into Equation (8) and writing it in a slightly different way as a summation over all subsets of size *i*, times an indicator of set inclusion. The fourth equality follows from the fact that there are exactly 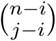 sets of size *i* that contain a particular set of size *j*. This follows from the fact that, given that the *j* specific elements are in the set of size *i*, there are 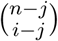 ways to choose the other *i*−*j* elements of the set. The fifth equality follows from the definition of ***ξ***_*i*_ in Equation (9) and the final equality follows from rearranging the identity 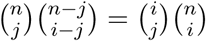.

If we define ***U***_*n*_ = (*U*_1_, …, *U* _*n*−1_) and ***ξ***_*n*_ = (*ξ*_*n*,1_, …, *ξ*_*n,n*−1_), we can rewrite Equation (11) in matrix form to yield the particularly simple matrix relationship

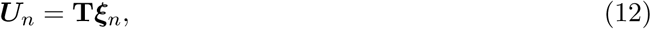

where **T** is the lower triangular matrix of dimension (*n* − 1) × (*n* − 1) whose element (*i, j*) is given by

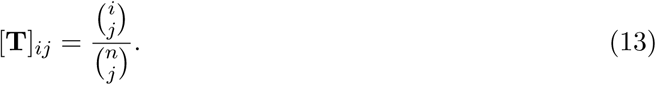

Because **T** is a lower-triangular matrix with non-zero diagonal elements (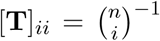 on the diagonal), its determinant is nonzero and it is therefore invertible. As we will show in the more general case of multiple populations in Section 3.5, the inverse transformation from ***U*** to ***ξ*** in one population can be expressed as

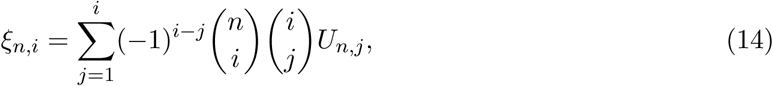

which has the more compact matrix representation ***ξ*** = **T**^−1^***U*** where **T**^−1^ is the lower triangular matrix whose elements are given by

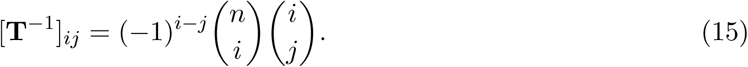

### 3.2. Computing the expected number of private segregating sites in a single population

We now obtain an expression for 𝔼*U*_*n,i*_ = 𝔼*π*(*S*_*i*_) in a single population by computing the expected number of sites that are private to a subset *S*_*i*_. Suppose that *n* sequences are sampled at time *t* = 0 (the present) and let *𝒜* _*n*_(*t*) denote the random number of ancestral lineages remaining at time *t* in the past. Let *L*_*n*_(*r, s*) denote the sum of total branch lengths in the genealogy between times *r* and *s* in the past with *r* ≤ *s*. Then *L*_*n*_(*r, s*) can be expressed as an integral over the expected number of ancestral lineages:

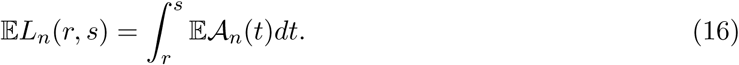

The result in Equation (16) was stated as an asymptotic approximation by Chen and Chen (2013) in the limit as *n* → ∞ (Equation 29 of that paper) and it was proved to hold exactly for finite n by Jewett and Rosenberg (2014) (Theorem 2.1 of that paper). Using Equation (16), Jewett and Rosenberg showed that under the infinite sites model, the expected total number of mutations *d*_[*r,s*]_(*S*_*n*_) private to a set *S*_*n*_ of *n* homologous sequences arising during the time interval [*r, s*] is given by

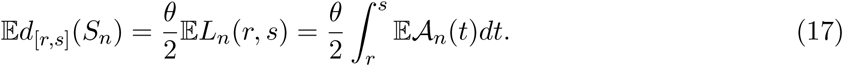

Equation (17) can be used to compute the expected number of sites 𝔼*π*(*S*_*i*_) that are private to a subset of *i* sampled sequences. In particular, if *d*_[*r,s*]_(*S*_*n*_) is the total number of mutations private to the full sample of size *n* in the time interval [*r, s*] and *d*_[*r,s*]_(*S*_*n*_\*S*_*i*_) is the number of mutations private to the set *S*_*n*_\*S*_*i*_ of *n* − *i* other sequences, then

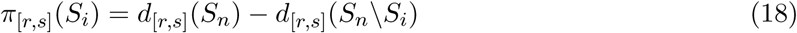

is the number of mutations arising in the time interval [*r, s*] that are private to the sequences *S*_*i*_, and no others. Taking the expectation of both sides and invoking Equation (17) gives

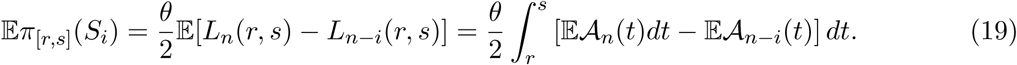

Equation (19) provides a simple way to compute expected numbers of private sites by integrating over expected numbers of ancestral lineages.

### 3.3. Computing 𝔼*ξ*_*n*_ exactly in a single population

Equation (19) gives us a way to compute *U*_*n,i*_, (and hence ***ξ***_*n*_ using Equation 14) as long as we can compute the expected number of ancestors 𝔼*𝒜*_*n*_(*t*) as a function of time. In a population with time-varying relative size *ν*(*t*), the expected number of ancestors can be computed exactly using the following expression due to Tavaré (1984) (Eqn. 5.11):

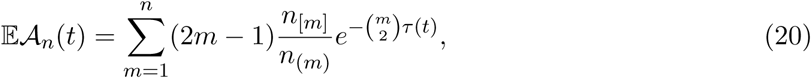

where *τ*(*t*) is the scaled coalescence time given by

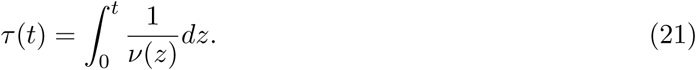

In Equation (20), the quantities *n*_[*m*]_ = *n* (*n* − 1) … (*n* − *m* + 1) and *n* _(*m*)_ = *n* (*n* + 1) … (*n* + *m* − 1) are the *m*th falling and rising factorials of *n*. If the population has constant size *ν*(*r*) in the time interval [*r, s*], then integrating both sides of Equation (20) gives

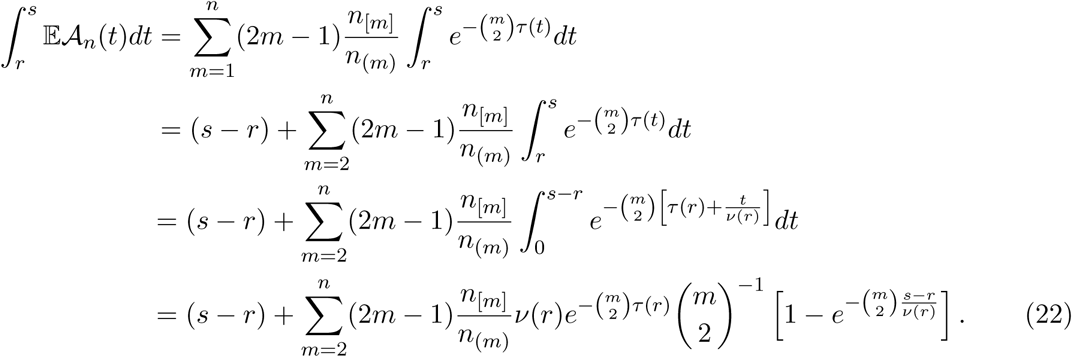

Combining Equation (22) with Equation (19) gives the expected number of private sites in a set of *i* sequences arising in the time interval [*r, s*] in which the population has constant relative size *ν*(*r*):

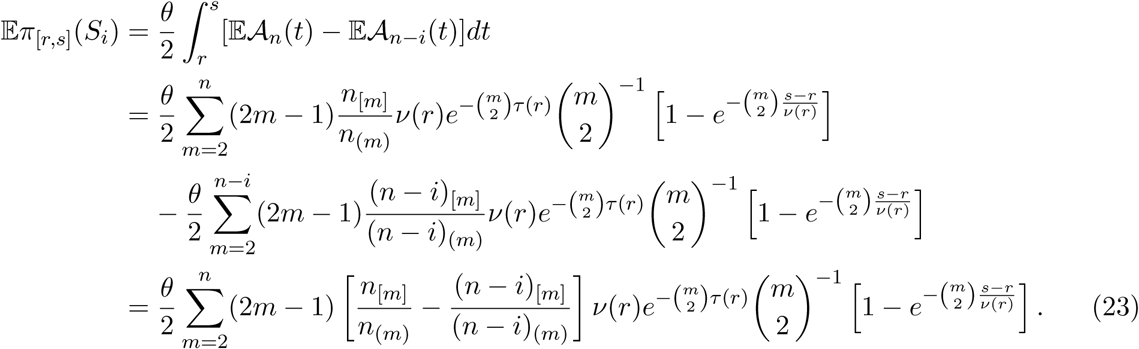

In Equation (23), we were able to combine the two summations in the second equality because (*n* − *i*)_[*m*]_/(*n* − *i*)_(*m*)_ = 0 for *m* > *n* − *i*; thus, the upper bound in the second summation can be set to *n*.

For a population history that is piecewise constant with relative size *ν*(*t*_*k*−1_) in each of the *K* epochs 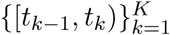, summing over all time intervals gives

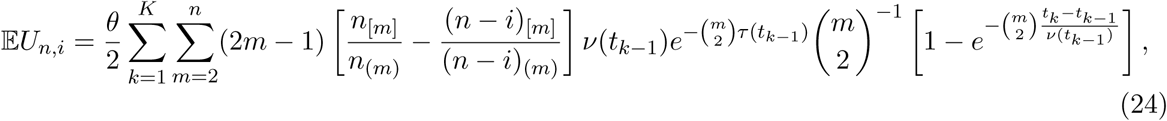

where 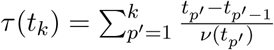. Finally, taking the expectation of both sides of Equation (14) and plugging in Equation (24) gives

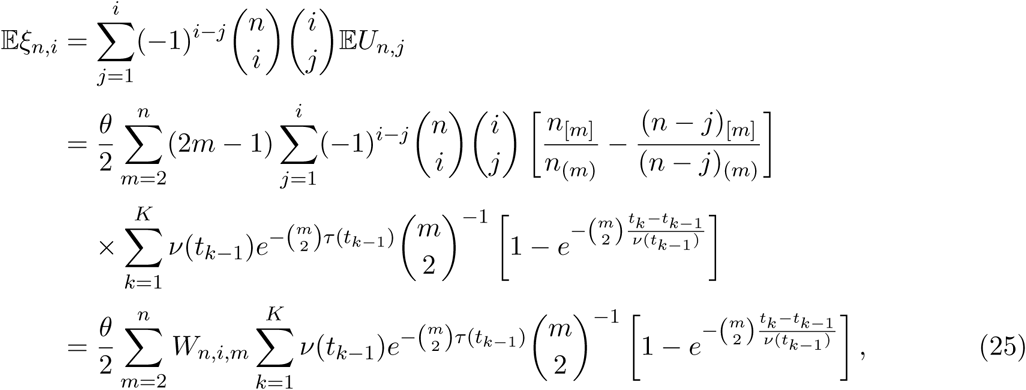

where

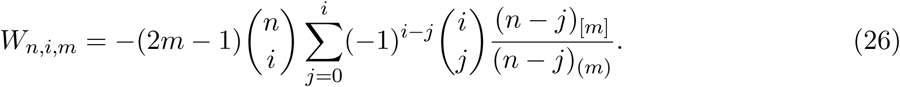

In Equation (26), we have set the lower bound on the summation to *j* = 0 because *n*_[*m*]_/ *n* _(*m*)_ − (*n* − *j*)_[*m*]_/(*n* − *j*)_(*m*)_ = 0 when *j* = 0. Moreover, the term *n* _[*m*]_/ *n* _(*m*)_ in the second equality of Equation (25) drops out because it is constant in *j* and 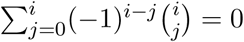 by the binomial theorem.

By comparison with Equation (10) of Kamm et al. (2017), it can be seen that the two formulas are the same. In particular, the quantity *W*_*n,i,m*_ in Equation (26) is precisely the quantity *W*_*n,i,m*_ in Kamm et al. (2017) and the internal summation over *k* in Equation (25) is the explicit form of the integral 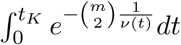 for a population of constant size. Our approach provides a closed formula for the term *W*_*n,i,m*_, which was found by a recursion in Kamm et al. (2017), following the derivation in Polanski and Kimmel (2003). However, because the recursion is faster to compute in practice, we provide the recursive form of *W*_*n,i,m*_ in the final presentation of the formula in Box 1.

### 3.4. Approximating 𝔼*ξ*_*n*_ in a single population

So far, we have computed exact formulas for the SFS. However, in preparation for deriving approximate formulas for the SFS that are computationally efficient in the case of multiple populations with pulse migration events, we first consider the approximation in a single population.

An approximate expression for *ξ*_*n,i*_ can be obtained by following the the same approach used in Section 3.3, but replacing the exact formula for 𝔼*𝒜* _*n*_(*t*) (Equation 20) with an approximate formula. The simplicity and computational efficiency of existing approximations of 𝔼*𝒜* _*n*_(*t*) make it possible to obtain fast approximate formulas for 𝔼*π*_[*r,s*]_(*S*_*i*_), allowing us to obtain fast approximations of the SFS.

In a single population, Griffiths (1984) showed that the expected number of ancestors at time *t* in a population with relative size *ν*(*t*) for *t* ∈ [0, ∞) can be approximated by

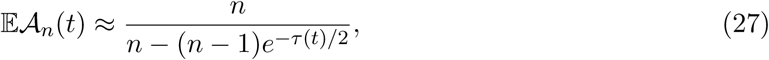

where *τ*(*t*) is the scaled coalescence time given in Equation (21). Griffiths showed that the approximation in Equation (27) holds asymptotically as *n* → ∞ or as *t* → 0.

If the population has constant size in the time interval [*r, s*], then integrating both sides of Equation (27) over the time interval [*r, s*] gives (Appendix C)

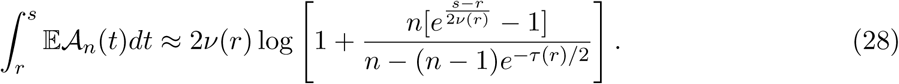

If the population has piecewise constant size defined over the intervals 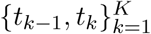 in which *ν*(*t*) = *ν*(*t*_*k*−1_) for *t ∈* [*t*_*k*−1_, *t*_*k*_) and *t*_0_ < *t*_1_ < … < *t*_*K*_ < ∞, then by combining Equation (28) with Equation (19), we find that the number of sites private to *i* sequences that arise in the *k*th time interval is given by

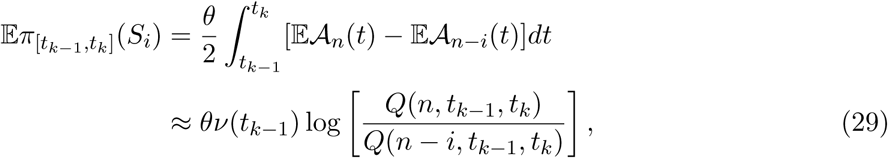

where *Q*(*n, r, s*) is given by

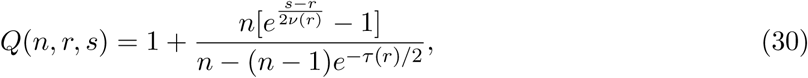

and the scaled time in Equation (30) is given by

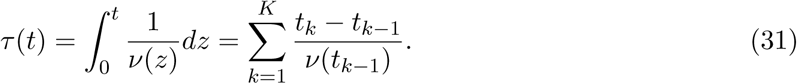

Summing Equation (29) over all time intervals and using the fact that 𝔼*U*_*n,i*_ = 𝔼*π*(*S*_*i*_) (Equation 8) gives

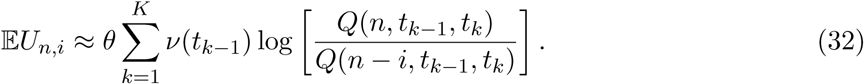

Finally, plugging Equation (32) into the expectation of Equation (14) gives

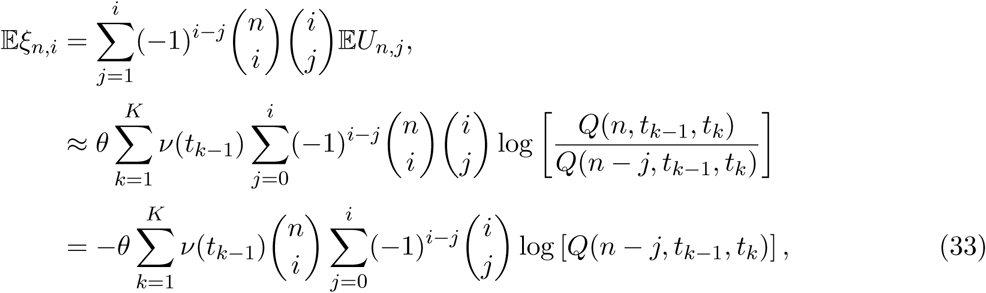

where the lower bound on the summation in the second equality can be taken to zero because the summand is zero at *j* = 0 and the final equality follows from the fact that *Q*(*n, r, s*) is constant in *j* and 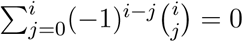 by the binomial theorem. This gives the approximation of *ξ*_*n,i*_ in Equation (3) of Box 1.

### 3.5. The relationship between *U*_n_ and *ξ*_n_ in multiple populations

The derivation of the relationship between ***ξ***_**n**_ and ***U***_**n**_ in multiple populations follows the same approach as the derivation in the case of a single population. In the case of *P* different populations with samples of sizes *n*_1_, …, *n* _*P*_, respectively, let 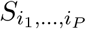 denote a subsample of sequences with *i*_*p*_ lineages in population *p*, for *p* = 1, …, *P*. As in the case of a single population, define 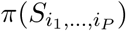 to be the number of private sites in the sample and define 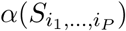 to be the number of sites that are ancestral to the subsample and to no other sequences.

The multi-population forms of 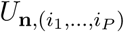 and 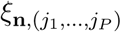, are

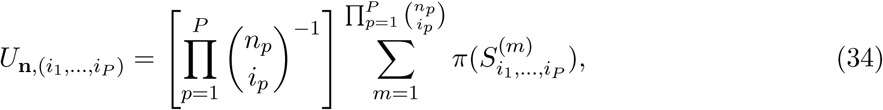

and

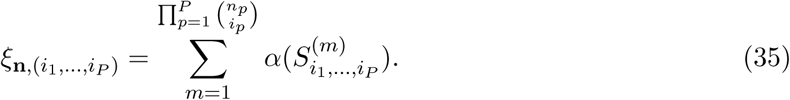

The relationship between 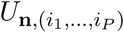 and 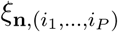 can be established by the same approach we used to derive Equation (11). This derivation is carried out in Appendix A and gives

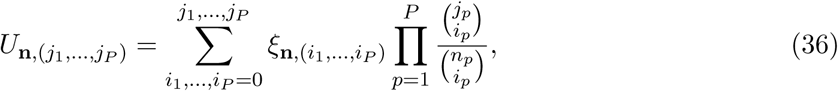

where we take ***U***_**n**,(0,…,0)_ = ***ξ***_**n**,(0,…,0)_ = 0. It is straightforward to check that the inverse transformation from ***U***_**n**_ to ***ξ***_**n**_ is given by

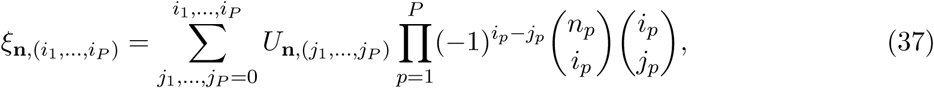

which can be checked by plugging Equation (37) into Equation (36) and showing that the composite transformation yields the identity. We carry out these calculations in Appendix B.

### 3.6. Approximating *U*_n_ in multiple populations with piecewise constant sizes and pulse migrations

The results of Sections 3.2 through 3.5 can be combined to obtain a fast approximate formula for ***ξ***_**n**_ in a collection of piecewise constant populations connected by pulse migrations. In particular, for a set of *P* populations with *n*_*p*_ lineages sampled from population *p*, for *p* = 1, …, *P*, let *𝒜* _**n**_(*t*) denote the total number of ancestors at time *t* of the set 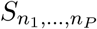 of *n*_1_, …, *n*_*P*_ sampled sequences. Similarly, let *𝒜* _**i**_(*t*) denote the total number of ancestors at time *t* of a subset 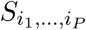 of *i*_1_, …, *i*_*P*_ sequences. Finally, let *𝒜* _**n**,*p*_(*t*) denote the number of ancestors of *n*_1_, …, *n*_*P*_ sequences that exist in population *p* at time *t*, and similarly define *𝒜* _**i**,*p*_(*t*).

If the size of each of the *P* populations is constant in a time interval [*r, s*], then the total number of alleles arising in the time interval [*r, s*] that are private to the subsample 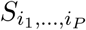 can be found using Equation (19) as

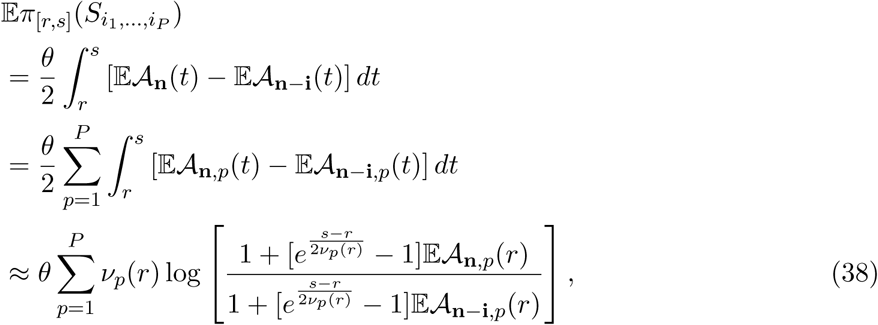

where the final equality comes from Equation (C.5) in the Appendix.

Now suppose that the relative size of the pth population is *ν*_*p*_(*t*) = *ν*_*p*_(*t*_*k*−1_) in the time interval *t ∈* [*t* _*k*−1_, *t*_*k*_), for *K* different time intervals 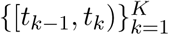 satisfying *t*_0_ < … < *t*_*K*_ < ∞. Then the total number of private segregating sites over all time intervals is

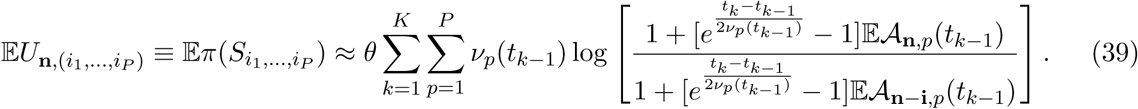

To find 𝔼*𝒜*_**n**,*p*_(*t*), we can use the fact that pulse migrations simply transfer ancestral lineages from one population to another at fixed times in the past. Let *m*_*k,p,p*′_ be the fraction of population *p*′ that instantaneously immigrates from population *p* at the top of interval *k*, looking forward in time and let ***M***_*k*_ be the *P* × *P* matrix whose (*p, p*′) entry is [***M***_*k*_]_*p,p*′_ = *m*_*k,p,p*′_ if *p* ≠ *p*′ and 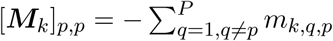. Given that 𝔼*𝒜* _**n**,*p*_(*t*_*k*−1_) ancestors exist in population *p* at the bottom of the *k*th time interval, then using Equation (27), the number 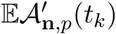 remaining at the top of the interval immediately before migration is approximately

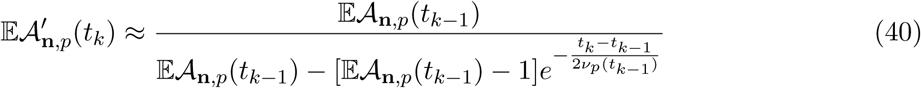

and the number remaining at the top after migration is

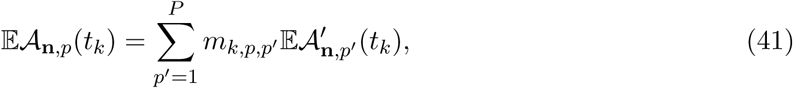

which has the more compact matrix representation

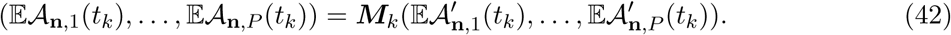

Thus, the quantities 𝔼*𝒜* _**n**,*p*_(*t*_*k*_) can be found by recursively applying Equations (40) and (42). Applying the relationship between 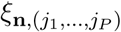 and 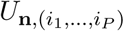 given in Equation (37) to Equation (39) gives the result in Equation (4) of Box 2.

## 4. Discussion

We have obtained accurate approximate formulas for computing the SFS in populations of piecewise constant size with instantaneous pulse migrations among them. The computational complexity of these formulas depends the index of the term of the SFS being computed, rather than on the sample size, allowing low-order terms of the SFS to be computed quickly for arbitrarily large sample sizes.

Our formulas for the SFS were derived by conceptualizing the SFS as a weighted sum over expected total ancestral branch lengths among *i* sampled lineages. In contrast, previous approaches expressed the SFS as a weighted sum over expected first coalescence times among 2, …, *n* lineages (Polanski and Kimmel 2003, Chen and Chen 2013, Kamm et al. 2017). Conceptualizing the SFS in terms of expected sums of branch lengths makes it possible to obtain the simple and fast formulas we derive here. The approach is useful more generally for deriving approximate coalescent formulas under complicated demographic models (Jewett and Rosenberg 2014).

It is important to note that approaches based on the diffusion approximation of the Wright-Fisher model are still the most efficient methods for computing the full SFS when the number of populations is small. However, computing the full SFS becomes intractably slow as the number of samples and populations increases. Thus, the benefit of the approximate formulas we present is their higher efficiency for computing a subset of SFS terms when sample sizes are large. The approximations derived here are also more accurate than the diffusion approximation for low order terms of the SFS when the number of sampled haplotypes is moderate or large.

The formulas we obtain are for populations of piecewise constant size with instantaneous pulse migrations. However, they can be used to approximate the SFS for populations of time-varying size and continuous migration by taking the time-step to be short. It is also possible to derive approximations of the SFS in the case of exponentially growing populations and continuous migration by substituting approximate or exact formulas for the expected number of ancestral lineages under these scenarios into the penultimate equality of Equation (38). Approximations for the expected number of ancestral lineages under continuous migration and arbitrary size changes are given in Jewett and Rosenberg (2014). However, for populations with exponentially changing sizes the approach described here yields formulas that are computationally less efficient and numerically less stable than existing methods. Thus, we have chosen to focus on the fast approximations for piecewise constant populations presented in this paper.

## 5. Acknowledgments

This research was supported in part by a National Institutes of Health grants R01-GM109454 and R01-GM094402 and by a Packard Fellowship for Science and Engineering awarded to Yun S. Song. I am very grateful to Yun Song for his support and multiple close readings of the manuscript. Many thanks also to Jeffrey Spence for lots of helpful discussions and suggestions.

## Appendix A. The relationship between 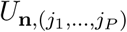 and 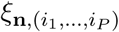

To derive the relationship between 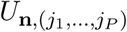 and 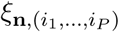, we begin with the definition of 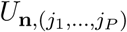:

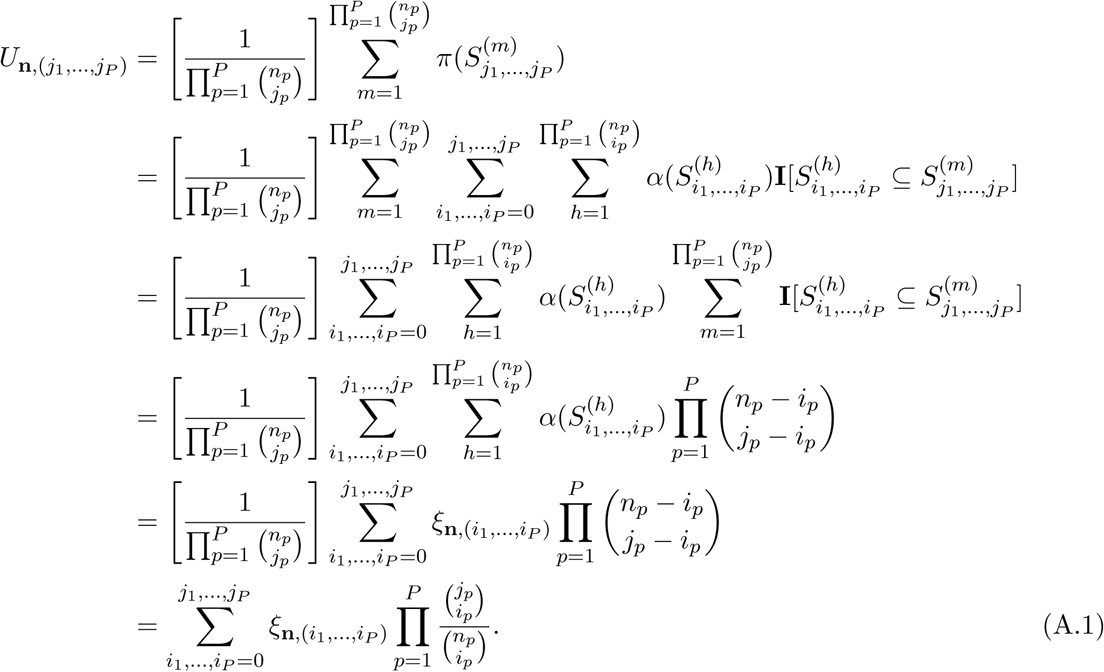

In Equation (A.1), the second equality follows from writing the summand as a sum over alleles ancestral to all subsets 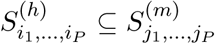 such that *i*_*p*_ ≤ *j*_*p*_ for *p* = 1, …, *P*. The fourth equality follows from the fact that 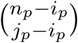 is the number of subsets of size *j*_*p*_ in population *p* that contain a particular subset of size *i*_*p*_. As in the single-population case, this result follows from the fact that there are 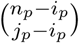 ways to choose the *j*_*p*_ − *i*_*p*_ other members of this subset. The fifth equality follows from the definition of 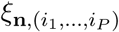 in Equation (35) and the final equality follows from rearranging the identity 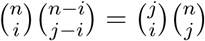.

## Appendix B. The inverse transform from 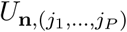 to 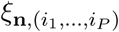

This transform can be derived by plugging Equation (37) into Equation (36) and showing that the composite transformation yields the identity:

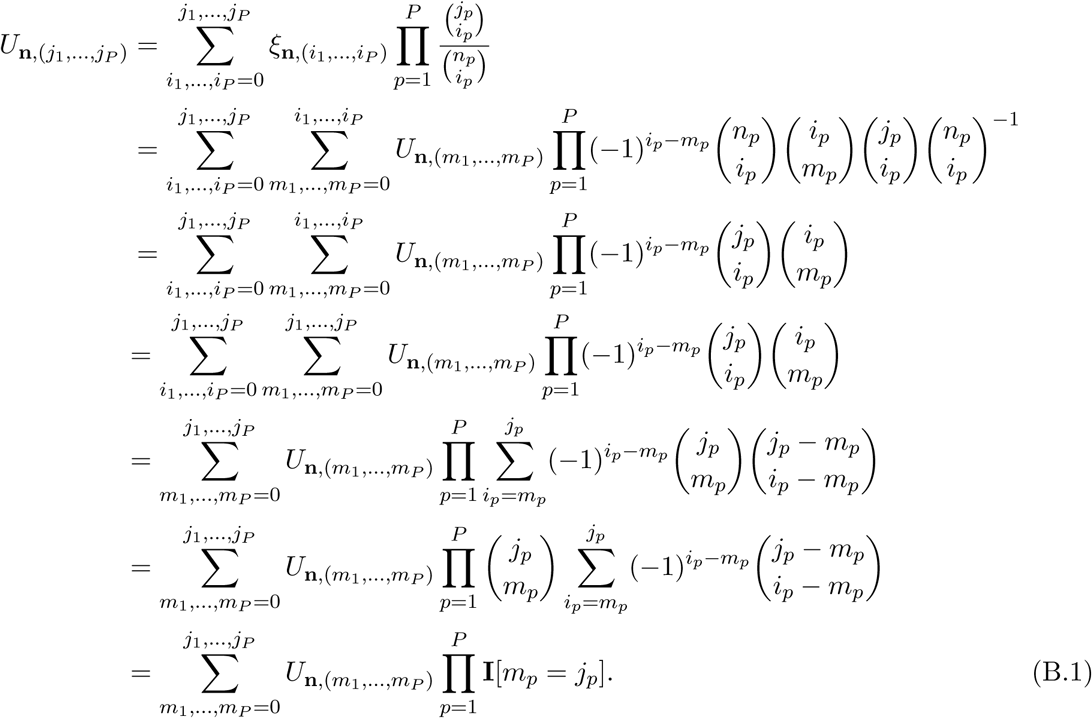

In the fourth equality, we have extended the upper bound of the inner summation up to *j*_1_, …, *j*_*P*_ using the fact that 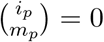 for *m* _*p*_ > *i* _*p*_. In the fifth equality, we brought the summation inside the product and used the identity 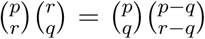. The final equality follows from reindexing *h*_*p*_ = *i*_*p*_ − *m*_*p*_ and noting that the summation 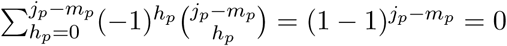 whenever *j*_*p*_ ≠ *m*_*p*_ by the binomial theorem, and it is equal to one if *j*_*p*_ = *m*_*p*_. Thus, we arrive at the fact that the right-hand side of Equation (B.1) is equal to 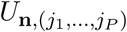, proving the identity.

## Appendix C. Approximation of 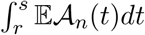

The derivation of Equation (28) amounts to a change of variables and some algebra. First, noting that *τ*(*t*) = *τ*(*r*) + (*t* − *r*)/*ν*(*r*) for *t ∈* [*r, s*] whenever the relative population size is constant in [*r, s*], we have

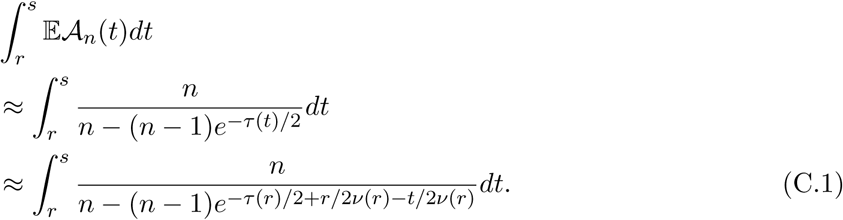

Thus, setting *b* = −(*n* − 1)*e*^−*τ*(*r*)*/*2+*r/*2*ν*(*r*)^, *c* = −1/2*ν*(*r*) we have an integral of the form

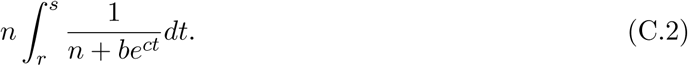

Making the change of variables *y* = *e*^*ct*^ so that 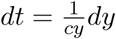, we have

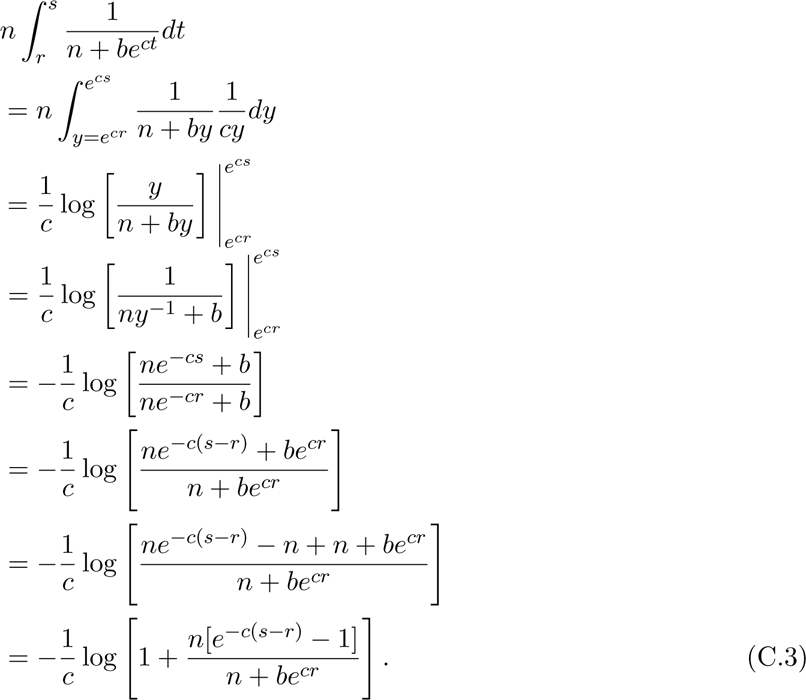

Plugging in *b* = −(*n* − 1)*e*^−*τ*(*r*)*/*2+*r/*2*ν*(*r*)^ and *c* = −1/2*ν*(*r*), we obtain the result in Equation (28):

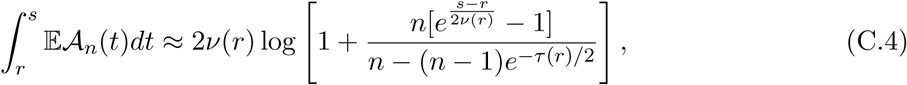

Noting that *n*/(*n* − (*n* − 1)*e*^−*τ*(*r*)*/*2^) *≡𝒜*_*n*_(*r*), we can express Equation (C.4) more compactly as

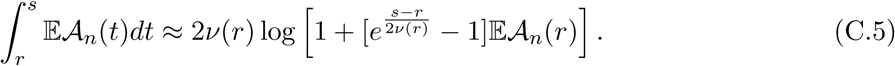

Equation (C.5) allows us to approximate the expected branch length in a time interval [*r, s*] as long as we know the number of ancestors remaining at time *r*.

